# Accounting for sources of bias and uncertainty in copy number-based statistical deconvolution of heterogeneous tumour samples

**DOI:** 10.1101/004655

**Authors:** Christopher Yau

## Abstract

Deconvolving heterogeneous tumour samples to identify constituent cell populations with differing copy number profiles using whole genome sequencing data is a challenging problem. Copy number calling algorithms have differential detection rates for different sizes and classes of copy number alterations. This paper describes how uncertainty in classification and differential detection rates can introduce biases in measures of clonal diversity. A simulation strategy is introduced that allows differential detection rates to be adjusted for and this process is shown to minimise bias.

## Background

Somatic copy number alterations (SCNAs) and genomic instability are a hallmark of many cancers [21]. These somatic changes can lead to the loss or gain of function where the underlying genes affected have a regular tumour suppressor or oncogenic promoting activity. Genome-wide identification of SCNAs in cancers has been enabled by the use of high-throughput genomic technologies including microarray-based comparative genomic hybridisation (aCGH), single nucleotide polymorphism (SNP) microarrays [1, 21] and, more recently, whole genome sequencing (WGS) [11, 9]. The focus of this article will be on whole genome sequencing analysis.

A wide variety of advanced bioinformatics tools have been developed to support these applications [12, 3, 4, 6, 20, 7, 10, 19, 13]. These tools tackle a number of computational and statistical modelling challenges involved in the analysis of heterogeneous cancer data. These include tackling technical issues (“noise”) such as variation due to local genomic GC content, sequence mapping errors, read errors, etc. Furthermore, the purity and ploidy of a tumour sample are typically unknown quantities and correctly estimating these parameters can have a substantial impact on the interpretation of the data.

Copy number analysis of heterogeneous tumour samples involves four tasks: (i) identification of genomic segments with homogeneous data features, (ii) classification of each segment with the type of copy number alteration involved, (iii) quantifying the prevalence of the copy number alteration (the proportion of cells in the tumour sample that exhibits each event) and (iv) determining the cellular constitution of the tumour sample (the number of different cell types and the copy number aberrations contained by each cell type). The first three of these tasks have received much attention but the investigation of tumour sample constitution is relatively under-developed and a topic of ongoing current research interest. Copy number calling differs from the identification of single nucleotide variants similarities as copy number aberrations affect genomic regions as supposed to specific loci. This introduces an additional spatial dimension to the problem.

Inferring tumour sample constitution is a challenging problem as current sequencing technologies and protocols are unable to determine the cell-of-origin for sequence reads (except in the special case of single cell sequencing where cells are physically separated [22, 18, 8]). Consequently, if a tumour is heterogeneous and contains a mixture of cell types, the observed sequence data only provides an aggregated measure of the combined effect of all the cell types in the sample. Success in the statistical deconvolution of heterogeneous tumour samples has predominantly arisen from the analysis of single nucleotide variants (SNVs) using high-depth targeted or exome sequencing in conjunction with computational methods [5, 16, 13]. For constituent populations that are separated by copy number aberrations, THetA (Tumor Heterogeneity Analysis) [10] was developed recently to infer the collection of constituent genomes and their proportions in a tumour sample using a maximum likelihood-based statistical deconvolution framework that considers read depth information from segmented tumour genomes. Deconvolved cancer genomes can be used to provide insights into tumour heterogeneity and the evolution of a tumour. The consequences and implications of genetic heterogeneity in cancer have received much interest recently [2, 15, 17].

Whilst computational methods have been widely developed to detect copy number aberrations from whole genome sequencing data, the ability to thoroughly validate computational investigations is being outpaced by the decreasing costs and widespread availability of high-throughput sequencing facilities. Complimentary techniques, such as fluorescence *in situ* hybridisation (FISH), are labour-intensive, costly and too low-throughput to comprehensively validate the many findings reported from WGS studies. State-of-the-art single cell techniques themselves suffer from technical limitations (e.g. whole genome amplification biases, allelic dropout) and therefore do not necessarily represent a gold standard for very high resolution analysis. This is in contrast to the validation of SNVs where Sanger sequencing, PCR and targeted sequencing can be used to easily and rapidly verify the existence of putative mutations. As a consequence, with copy number aberrations, there is an increased reliance on raw computational findings in comparison.

The purpose of this investigation is to consider this reliance rigorously. We ask how biases and uncertainties in the performance of a copy number calling algorithm could impact on downstream analyses with a particular focus on the identification of sub-clonal copy number aberrations and measures of clonal diversity that maybe derived from these findings. We will demonstrate that there are intrinsic detection limits in the identification of copy number aberrations and that failing to account for differential detection rates of a copy number caller can lead to inflated measures of clonal diversity. We then suggest a simulation strategy that can be adopted to account for this to produce robust measures of clonal diversity. The study provides a cautionary tale about the need for the propagation of uncertainty information in genomic data analysis especially where statistical tools might be built upon the output of another.

In order to explore the details of the deconvolution problem, an existing statistical modelling framework is utilised to illustrate the potential issues. The model integrates multi-scale genome segmentation, absolute allelic-specific copy number classification, ploidy estimation and the computation of the prevalence of copy number events in heterogeneous tumours. These statistical methods are implemented in a software package called OncoSNP-SEQ2, an update of an earlier version of the software [19]. However, it should be stressed that the purpose of this study is not to evaluate the relative performance of the copy calling method itself but to highlight the processes that might be involved in determining how the characteristics of a chosen copy number caller might impact on downstream analyses. These issues are intrinsic to the scientific problem and would be universally applicable to other methods.

## Results

### Methods Overview

Figure 1 shows an example genome-wide copy number analysis provided by OncoSNP-SEQ2. OncoSNP-SEQ2 adopts a multi-scale segmentation strategy to generate a series of copy number profiles of increasing resolution. Each series is associated with a rank level. Low ranked segmentations are produced using high penalty scores on segments thus resulting in “smoother” segmentations and generally contain larger SCNAs affecting large or whole chromosomal segments. Higher ranked segmentations are generated using lower penalty scores and contain increasingly smaller aberrations. Copy number calls produced by OncoSNP-SEQ2 consist of a set of segments where, for each segment, the total and minor copy number are reported as well as the prevalence of the SCNA in increments of 10%. Each call is associated with the rank of the segmentation in which that particular SCNA first arose. Later, in the simulation studies, we will filter calls by their ranks to vary the false positive detection rate of the calling method. Further details of OncoSNP-SEQ2 are provided in Materials and Methods.

**Figure 1:**
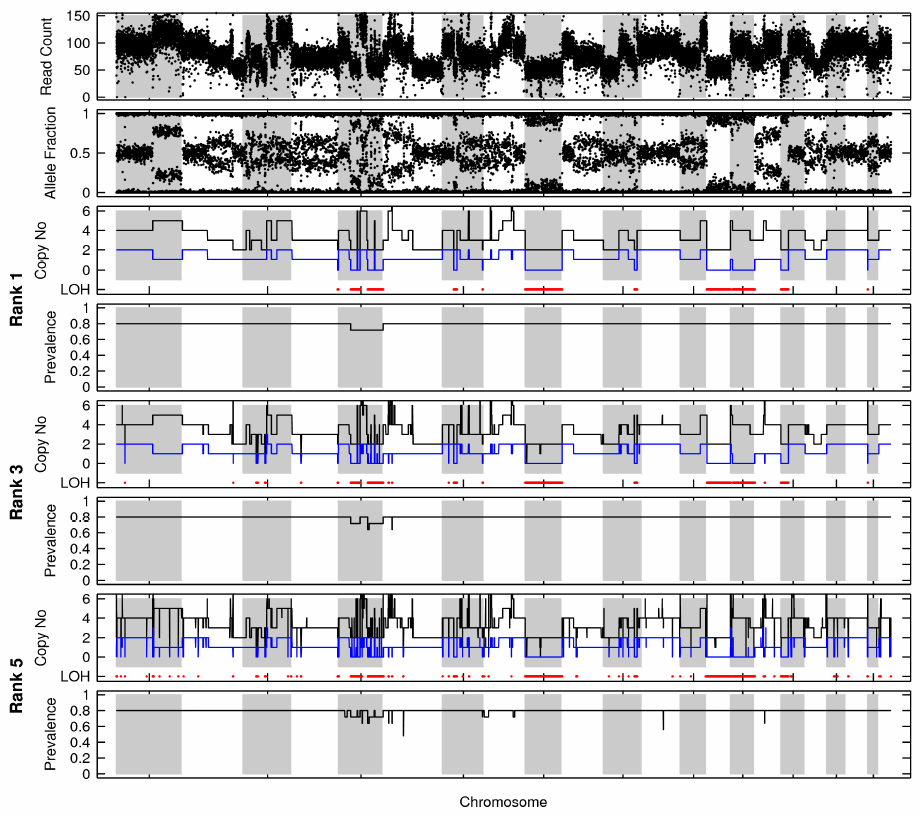
Example Copy Number Analysis. (a) Genome-wide read coverage and allele fractions, (b) probability map showing log-likelihood of different normal contamination and haploid coverage levels with three peak values identified and ordered by likelihood value and (c) low, medium and high-resolution copy number segmentations (black - total copy number, blue - minor copy number, red-loss-of-heterozygosity).

### Power to detect sub-clonal aberrations

We first performed a study to establish the differential detection rates of OncoSNP-SEQ2 using simulated datasets containing sub-clonal events of various sizes and prevalences. Whole genome sequencing data series (approximately 60X) was obtained from a chronic lymphocytic leukemia patient (CLL077) sequenced as part of a whole genome sequencing study [14] and allele-specific count data was obtained from approximately 1M SNPs. The sample series contains three substantial sub-clonal copy neutral LOH (CN-LOH) events on chromosomes 19 and 20 that were visually verified. These CN-LOH events were present in approximately (A) 30, (B) 60% and (C) 90% of cells respectively (see Figure 2(a-c)) and read counts for SNPs contained within these regions were extracted.

**Figure 2:**
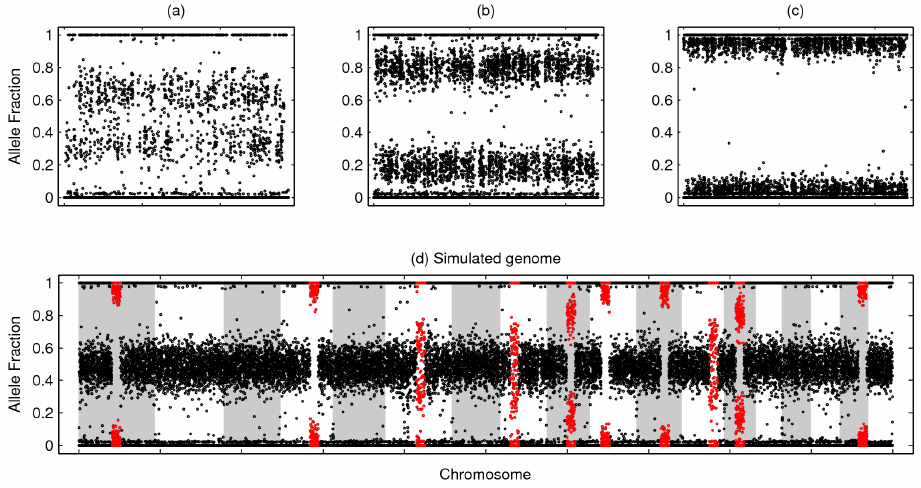
Simulated genomes. (a-c) Three CN-LOH events identified in patient CLL077. (d) Simulated genome containing randomly inserted CN-LOH events resampled from the extracted events (red). Sizes of inserted events are exaggerated here for the purposes of graphical illustration.

Next, three sets of 100 simulated tumours were generated. Each set of tumours contained the original CLL077 data from chromosomes 1-18 along with ten inserted CN-LOH events of various sizes obtained by resampling SNP data from one of the three verified CN-LOH events. Resampling from real events gave us the best chance of mimicking real-life observations of actual structural alterations. OncoSNP-SEQ2 was used to analyse each simulated data set. The true positive detection rate was measured as the average proportion of inserted CN-LOH events identified. An event was considered correctly classified if, at least, 50% of SNPs in the region were correctly called. The false positive rate was computed as the average proportion of SNPs, outside of the inserted CN-LOH regions, that were called as copy number alterations or LOH events.

Figure 3 shows that the power to detect sub-clonal events depends quite critically on the size of the CN-LOH event, its prevalence and the resolution of the copy number analysis. CN-LOH events that were larger and more prevalent in the tumour samples were more frequently detected and correctly classified even when using low ranked calls that produced low false positive rates. In contrast, smaller and/or less prevalent CN-LOH events could only be identified if much higher false positive rates are tolerated using higher ranked segmentations. The CN-LOH events at 30% prevalence were virtually undetectable unless they spanned five thousand SNPs.

**Figure 3:** Detection Rates. Detection results for copy-neutral LOH events of various lengths (*d*) used in the simulated data sets. LOH events had prevalences of (yellow) 90%, (red) 60% and (black) 30% respectively.

These results are unsurprising since small, low-prevalence sub-clonal CN-LOH events are difficult to distinguish from random signal fluctuations that commonly arise in current whole genome sequencing data due to technical sequencing artefacts. Furthermore, CN-LOH events can only be identified through changes in allele fractions at SNPs and not variation in read depth. At 60X, there is considerable variability in allele fractions and a large number of data points are required to distinguish a low-prevalence CN-LOH region from a normal region of heterozygosity. Higher depth sequencing would improve detection rates but the cost of current technologies means it is unreasonable to sample large numbers of tumours using coverages much above the 60X used in our simulations.

### Bias in clonal diversity estimates

After establishing that there are clear differential detection rates in the calling algorithm, we questioned whether the use of high resolution copy number analyses could introduce a bias in measures of clonal diversity for this tumour. A simple measure of clonal diversity that is applicable here would be the number of distinct prevalence states identified in the tumour sample as this would indicate the presence of multiple sub-populations in the sample. We tested this by examining a set of 100 simulated tumours where each tumour contained a mixture of the three CN-LOH event types. We analysed each simulated tumour sample with OncoSNP-SEQ2 and counted the number of distinct prevalence states detected. Counts were then stratified by the minimum event size and the rank of the segmentation in which the event was found.

Figure 4 shows that when using low-rank calls it was not always possible to identify the presence of all three CN-LOH event types in the simulated tumours. The identification of three distinct prevalence states was a relatively infrequent occurrence and it was more common to find just one or two of the prevalence states. It was also common to fail to detect any sub-clonal abnormalities. With higher ranked calls and allowing for smaller copy number aberrations it was possible to detect the existence of more prevalence states often above the three known to exist in the data. The increased number of prevalence levels gives an apparent appearance of high clonal diversity but this is likely to be purely artefactual. Relaxation of the stringency constraints has led to more than three distinct prevalence levels being identified in the tumour samples due to the inclusion of false positive calls. As previously established, power to detect small, low-prevalence CN-LOH events incurs a heavy price in greatly increased false positive rates.

**Figure 4:**
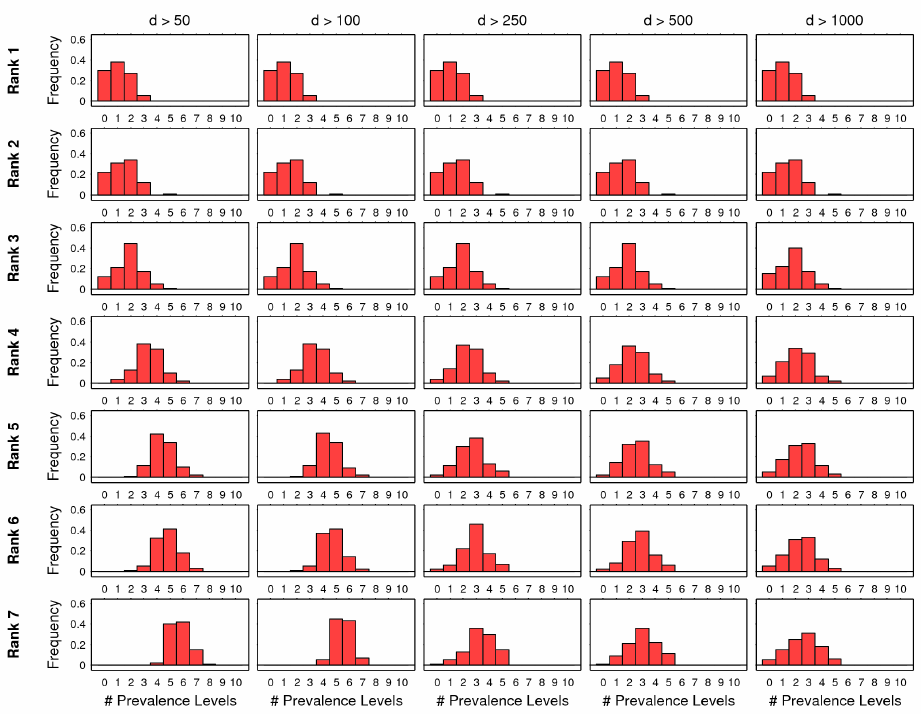
Prevalence Levels. Distribution of the number of detected prevalence levels across 100 simulated samples. Calls were stratified by rank (1 - Low resolution, 7 - High resolution) and minimum length of copy number event (*d*).

### Estimating differential detection rates

We next sought to determine if there are intrinsic limits in the detection rates of all classes of SCNAs besides CN-LOH. In order to answer this question, we conducted a simulation study to assess the recall accuracy for simulated SCNAs as a function of size, prevalence and class of aberration. As it is not practically possible to find real examples covering all possible SCNA classes and prevalence, our strategy was to generate blocks of data under the probability model used by OncoSNP-SEQ2. We then reclassified, under the same model (this would be the optimal classifier), to find if the original copy number class and prevalence used to generate the data could be recalled.

Specifically, total read depth and reference allele fraction data was simulated for a number of classes of SCNAs spanning blocks of varying number of SNPs and at different prevalence levels (see Table 2). For each simulated data block, the likelihood of the data was computed under each of the possible copy number classes and prevalence levels. If the most likely copy number class and prevalence matched the true values used to simulate the data matched then the class of the SCNA was said to have been successfully recalled. Note that in these simulations whole genome data was not simulated. Blocks of data corresponding to individual events were generated and classified without the need to do joint segmentation.

**Table 1:**
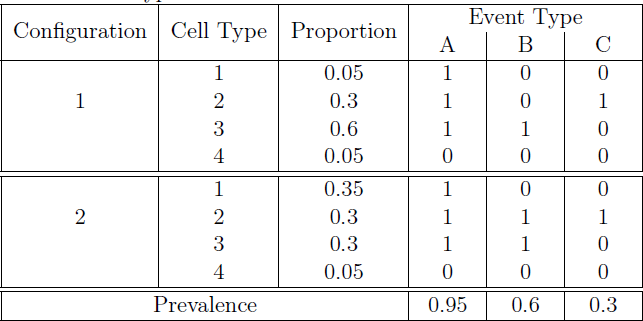
Multiplicity of sub-clonal architectures. Two alternate sub-clonal architectures that are compatible with the same observed cellular prevalences (0/1 indicates the absence/presence of an event in that cell type). Both configurations share two common cell types both differ in the third cell type.

| Configuration | Cell Type | Proportion | Event Type |  |  |
| --- | --- | --- | --- | --- | --- |
|  |  |  | A | B | C |
| 1 | 1 | 0.05 | 1 | 0 | 0 |
|  | 2 | 0.3 | 1 | 0 | 1 |
|  | 3 | 0.6 | 1 | 1 | 0 |
|  | 4 | 0.05 | 0 | 0 | 0 |
| 2 | 1 | 0.35 | 1 | 0 | 0 |
|  | 2 | 0.3 | 1 | 1 | 1 |
|  | 3 | 0.3 | 1 | 1 | 0 |
|  | 4 | 0.05 | 0 | 0 | 0 |
| Prevalence |  |  | 0.95 | 0.6 | 0.3 |

**Table 2:**
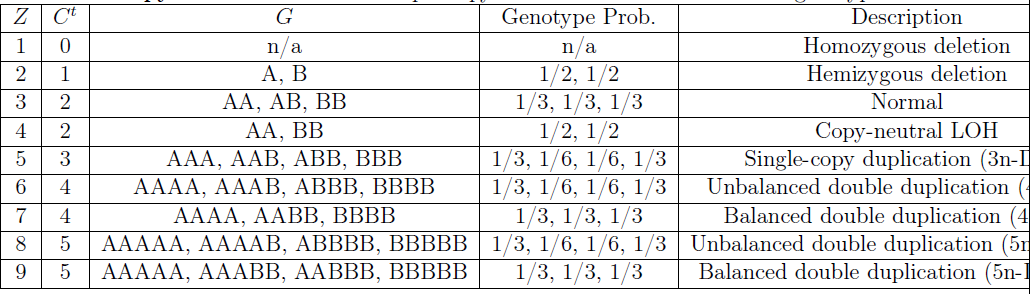
Copy number states. Example copy number states and associated genotypes.

Figure 5 shows the recall rates obtained via our simulation study for correcting recalling copy number only and for copy number and prevalence. Recall rates were near 100% for simulated SCNAs that were 100% prevalent (present in all cells of the tumour sample) and spanned hundreds of SNPs. However, it was more difficult to correctly classify *both* copy number state and prevalence than just to identify the copy number state alone. As the prevalence and/or size of the copy number event diminishes, recall rates fell and for most events there was a threshold size and/or prevalence at which recall accuracy degraded rapidly. For example, the copy number classes 4n-Dup and 5n-Dup (I) could not be recalled at below 50% prevalence. Note that at prevalences below 50% the expected data distribution for these classes overlap with those of the 3n-Dup class with high prevalence. Prior information in OncoSNP-SEQ2 is set to prefer higher prevalence states so the 3n-Dup class is preferentially chosen over the others.

**Figure 5:**
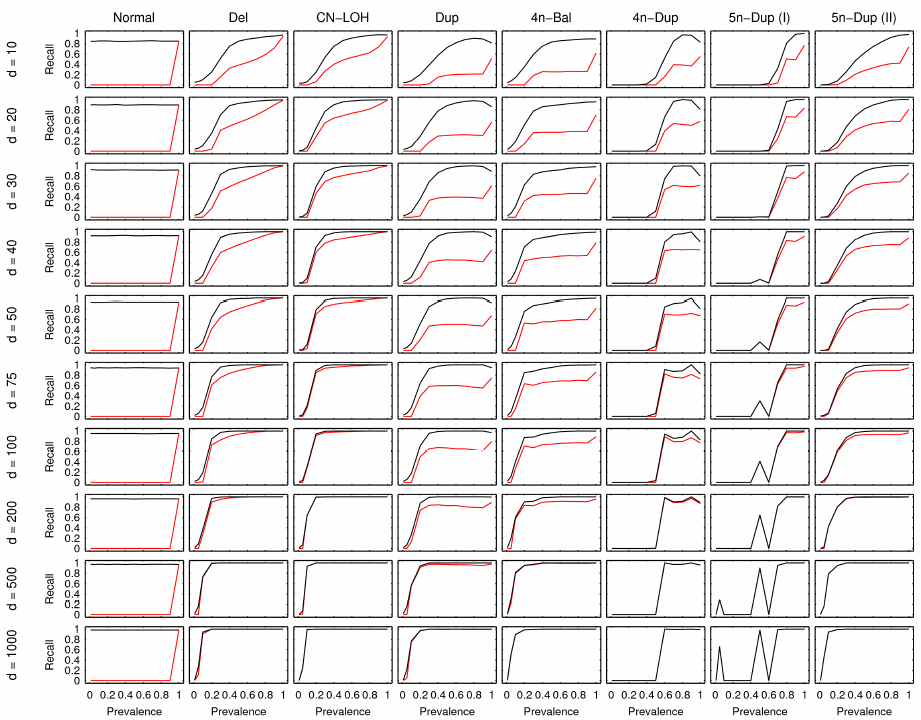
Theoretical Recall Rates. Plots showing optimal recall rates for different categories and sizes of copy number alterations at different prevalences. Recall rates are shown for (black) correct copy number recall and (red) for correct copy number *and* prevalence calls.

Some interesting oddities were also noted, for example, copy number events classed as 5n-Dup (I) (involving genotypes AAAAA, AAAAB, ABBBB, BBBBB) at prevalences of 0.6 had zero recall rate. This is because at this prevalence level the allelic fraction and expected read count for the 5n-Dup (I) class is nearly identical to that of a 4n-Dup class event (AAAA, AAAB, ABBB, BBBB), e.g. for the AAABB genotype with prevalence 0.6, the B allele fraction is given by (0.6 × 2 + 0.4 × 1)/(0.6 × 5 + 0.4 × 2) ≈ 0.26 and 0.6 × 5 + 0.4 × 2 = 3.8 times the haploid coverage which (when observed with noise) is close to the values of (0.25, 4) for a AAAB genotype.

### Adjusting for differential detection rates reduces bias in clonal diversity measures

The previous simulation results demonstrate that there are intrinsic limitations in detection rates. These limitations varied with the type of copy number class, size of event and its prevalence. With higher copy number classes, prior preferences towards high prevalence states could cause gross misclassifications. With this knowledge, we returned to the multiple sub-clone simulations and devised an approach to eliminate the false positive calls that were leading to strong biases in clonal diversity. The objective was to use the differential detection rates established in the previous section to select only those copy number classes and corresponding prevalence levels where there was high probability of successful recall. As a consequence, we removed all SCNA calls whose predicted type, size and prevalence had less than a 90% theoretical recall rate.

Figure 6 shows that when we account for the differential detection rates the number of estimated prevalence levels was substantially reduced even when high-ranked segmentations and small SCNA calls were used (contrast with Figure 4) whilst, at the same time, the number of simulated tumours with three detected prevalence levels (the true number) is increased. Therefore, by accounting for differential detection rates, it is possible to allow high resolution copy number analysis whilst minimising the occurrence of false positive calls that can inflate measures of clonal diversity. It is important to note that the filtering criteria are not based on arbitrary heuristics but optimally matched to the specific detection characteristics of the copy number caller.

**Figure 6:**
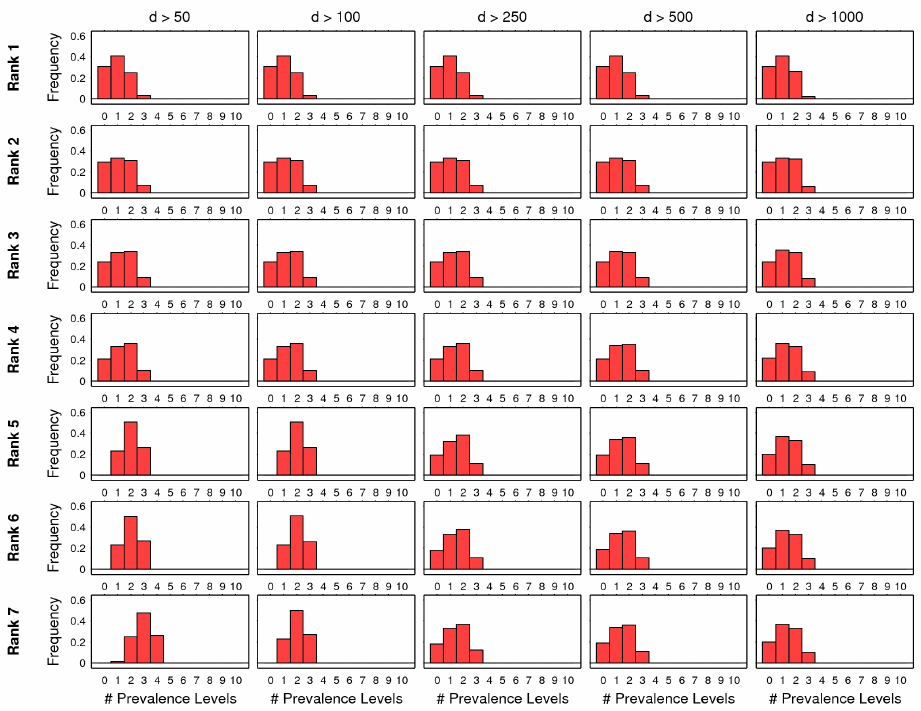
Adjusted Prevalence Levels. Distribution of the number of detected prevalence levels across 100 simulated samples adjusted for differential detection rates. Calls were stratified by rank (1 - Low resolution, 7 - High resolution) and minimum length of copy number event (*d*).

## Discussion

This study has shown that characterising the heterogeneous copy number landscape of a tumour requires careful thought. The signal processing necessary to detect copy number aberrations from whole genome sequencing data has certain intrinsic limits that are determined by the type, size and prevalence of the copy number aberrations.

It is a simple principle that a copy number caller cannot be relied upon to detect small, low-prevalence copy number alterations as reliably as large, whole chromosomal changes and therefore some account of these differential detection rates should be taken in downstream analyses. If all copy number calls are mistakenly treated as equally reliable then we have shown that there maybe negative consequences for measuring clonal diversity as false positive calls may drive up the apparent degree of heterogeneity in the tumour sample.

We have shown that, using a simulation-based strategy, it is possible to measure the detection characteristics of a copy number caller and correct for these potential biases. Although the strategy was only applied to OncoSNP-SEQ2 it is a generic approach that could be adapted to the particular features of any copy number calling method. This is a task that is best undertaken by individual development teams as different copy number callers can use the same raw sequence information in different ways. Whilst the focus here has been on SNP information other callers use other combinations of read-depth, split-reads, mate-pair, etc and it is not possible for us to assess all of these approaches.

Additionally, the focus of this article has been on the deconvolution of single tumour samples, but many of the issues extend to multi-sampling designs as well. Measures of clonal diversity based on copy number differences between tumour samples could be biased by false positive copy number calls due to differential detection rates. In this case sample-to-sample variability adds an additional dimension to be consider. The practices and strategies discussed in this study could also be applied to correct for these artefacts.

## Conclusion

This study has shown that the reliance on computational methods for dissecting tumour heterogeneity from whole genome sequencing data should be reinforced by physical validation techniques where possible (e.g single cell analysis or FISH). If computational findings are to be relied upon with out validation, differential detection rates by copy number calling algorithms should be measured and this information used to filter copy number calls in order to avoid bias in downstream analyses such as the assessment of clonal diversity.

## Materials and Methods

### Statistical Model Overview

Qualitative description of the model construction and further mathematical descriptions are given later in Materials and Methods. OncoSNP-SEQ2 uses two coupled hidden Markov processes: one to model the changes in copy number states at different loci along the genome and the other to denote the prevalence of the copy number alterations in the tumour sample. The use of Markov chains as a statistical model of the latent, unobserved copy number states is a standard approach for this problem, however, we believe that the use of a secondary chain to model the sub-clonal structure is a novel construction. This second chain has 10 states corresponding to prevalence levels 0, 10, 20,…, up to 90%. The use of dual-coupled Markov chains allows OncoSNP-SEQ2 to cluster data into similar copy number and prevalence classes whilst enforcing first-order spatial continuity.

#### Assumptions

We assume the presence of *K* distinct copy number profiles in a tumour sample that contains an unknown number of sub-clonal populations. The quantity *K* is an unknown. Note that *K* is not necessarily the number of distinct cell types since sub-clones may genetically differ in other ways, e.g. single nucleotide variants, small insertion-deletions, transactions, etc. However, for brevity, in the following discussion we ignore the distinction unless otherwise stated.

We will always assume that one of the *K* cell types corresponds to the normal, germline copy number profile. The model presented here can be naturally extended to model germline alterations by having a specific Markov chain for the germline copy number sequence but this adds unnecessary complexity and, for simplicity, we will always assume the germline cell type to be devoid of copy number alterations. Since many cancer sequencing studies will involve sequencing of normal-tumour pairs, distinguishing germline and somatic copy number changes can be effectively done via independent copy number analysis of the normal sample and removing or highlighting these changes when considering the tumour samples.

Finally, at each locus, we shall assume that only one type of somatic copy number alteration may exist amongst the *K* profiles. This limiting assumption disallows the presence of a deletion and duplication at the same location in different clones but greatly reduces the dimensionality of the problem to a tractable complexity level. Furthermore, as the sequencing data can measure the aggregated effect across all cell types, it would not be possible to predict the presence of distinct cell types with different copy number changes at the same locus from a single tumour sample.

#### Data

The data is assumed to be presented in a generic format consisting of two measurements at a series of SNPs: (i) the reference allele count and (ii) the total read count (optionally a standardised depth of coverage measure in non-overlapping windows centred on the SNP could be used in addition to the total SNP read count). In the following simulations we use the set of SNPs present on the Illumina Human Omni-Quad 1M SNP genotyping array but any high density set of SNPs with minor allele frequency greater than 5% will suffice (heterozygotes are desirable in order to add allele-specific information).

#### Modelling sub-clonal architecture

The derivation of our statistical model is illustrated in Figure 7. The genome-wide sequence of somatic copy number alterations is modelled as an *S*-valued Markov chain, *Z*. This sequence comprises the *union* of all the somatic copy number alterations across the *K* cell types because of our starting assumptions. In many standard HMM approaches, the copy number sequence *Z* is normally the primary and sole target of inference given observed sequence data *Y* and the Markov assumptions allow a first-order degree of spatial continuity to be enforced.

**Figure 7:**
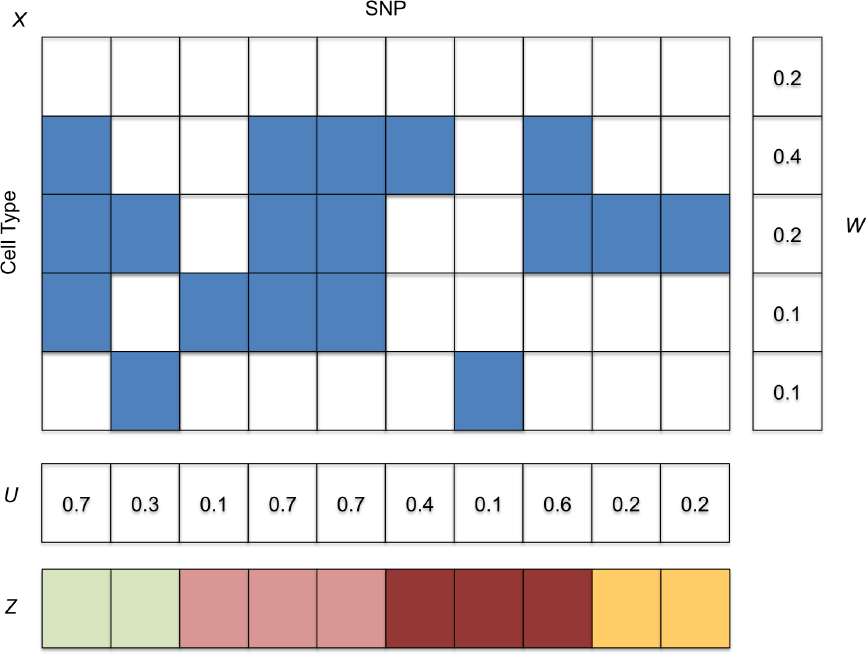
Model illustration. (Number of cell types, *K* = 5). A binary matrix is used to represent whether each cell type possesses a copy number alteration at a particular SNP. The proportion of the sample containing each cell type is given by a *K*-valued vector *W*. The copy number alteration type at each SNP are contained in a sequence *Z*. The sequence *U* represents the total proportion of the sample containing the copy number alteration. The observed sequence data can be explained with *Z* and *U* only and this is exploited in our model.

In this model, we wish to account for the fact that some cellular sub-populations may or may not feature the copy number alterations specified in *Z*. At each locus, given *Z*, each cell type can either possess a copy number alteration (feature) or not. Therefore the somatic copy number profile of the *K* cell types can be represented as a *K × N* binary feature allocation matrix, *X*, where *N* is the number of loci. In order to enforce first-order spatial continuity we specify that each row of *X* is modelled as a binary Markov chain. Each cell type is present in the tumour sample with proportions *W*_0_,…, *W*_*K*−1_. The first row of *X* is always zero as it corresponds to the germline profile and thus *W*_0_ has the special interpretation of being the degree of normal contamination.

Note that whilst this describes a possible model of the underlying sub-clonal architecture because current sequencing technologies do not provide cell-of-origin information for each sequence read, the expected allele-specific read counts and coverage levels are a function *only* of the genotype/copy number and the overall prevalence of the alteration at each location in the tumour sample. Hence, in order to explain the observed sequence data *Y*, the feature allocation matrix *X* is not directly required. It is only necessary to know the type of copy number alteration (*Z*) and the prevalence given by the sequence *U* = *XW*. We will discuss in the following why the quantity *U* is important for developing efficient, practical algorithmic implementations in the following.

#### Implementation issues

Typically, the number of populations *K* and their proportions are unknown *a priori* and are parameters to be inferred from the observed data. One formal (Bayesian) approach for comparing models of varying complexity is to compute the marginal likelihood of each model Pr(*Y |K* = *k*), *k* = 1,…, *K_max_* whilst integrating out uncertainty over all unknown parameters. This is computationally achievable in our model for small values, e.g. *K_max_* = 4 by using low-dimensional quadrature to marginalise over the weights *W* and using dynamic programming algorithms to exactly compute the marginal likelihoods Pr(*Y |W, K*) by enumerating all paths over the Markov structure. However, this approach is computationally demanding and becomes rapidly prohibitive as the model complexity increases exponentially with the number of cell types.

In order to develop a more tractable approach, we recognise that the observed data *Y* is only a function of *Z* and the sequence *U* = *XW*. As *X* is a binary feature matrix and each row is modelled as a Markov chain, the sequence *U* is also a Markov chain with 2*^K^* states corresponding to all the possible feature allocation combinations of *W* (e.g. for *K* = 3, the states are 0, *W*_0_, *W*_1_, *W*_2_, *W*_0_ + *W*_1_, *W*_0_ + *W*_2_, *W*_1_ + *W*_2_, *W*_0_ + *W*_1_ + *W*_2_ = 1). The exponential growth in the state space as a function of the number of cell types *K* is precisely why exact enumeration is computationally intractable for all except very small values of *K*. However, this observation that *U* forms a Markov chain allows us to develop an approximate algorithm whose computational costs are invariant to the number of cell types.

The principle underlying this method is to approximate the marginal distribution of *U* using a Markov chain with a dense, but fixed, state space, for example, with ten states corresponding to prevalences 0, 0.1, 0.2,…, 0.9. Statistical inference can then be conducted for (*Z, U*) only since knowledge of these is sufficient to fit the model to the observed data. Values of (*X, K, W*) can be *retrospectively* inferred from the posterior assignments to these states if desired. In essence we apply a two-level clustering approach that clusters loci into both discrete copy number *and* prevalence states. This model differs from the previous incarnation of OncoSNPSEQ [19] where the prevalence levels were treated as independent random variables and hence a prevalence level could not be directly associated with any SCNA.

### Prior calibration

OncoSNP-SEQ2 method computes the probability of the data under different combinations of haploid coverage and normal cell contamination values (see Figure 8). The haploid coverage and the degree of normal cell contamination determine the expected number of reads associated with a unit copy number change and hence their values directly impact the absolute copy number estimates. Local modes in the probability distribution over these parameters are automatically identified and, for each mode, a genome-wide copy number profile is reported. For many complex tumours, it is common to find multiple local modes in the probability distribution each corresponding to a different interpretation of the sequencing data.

**Figure 8:**
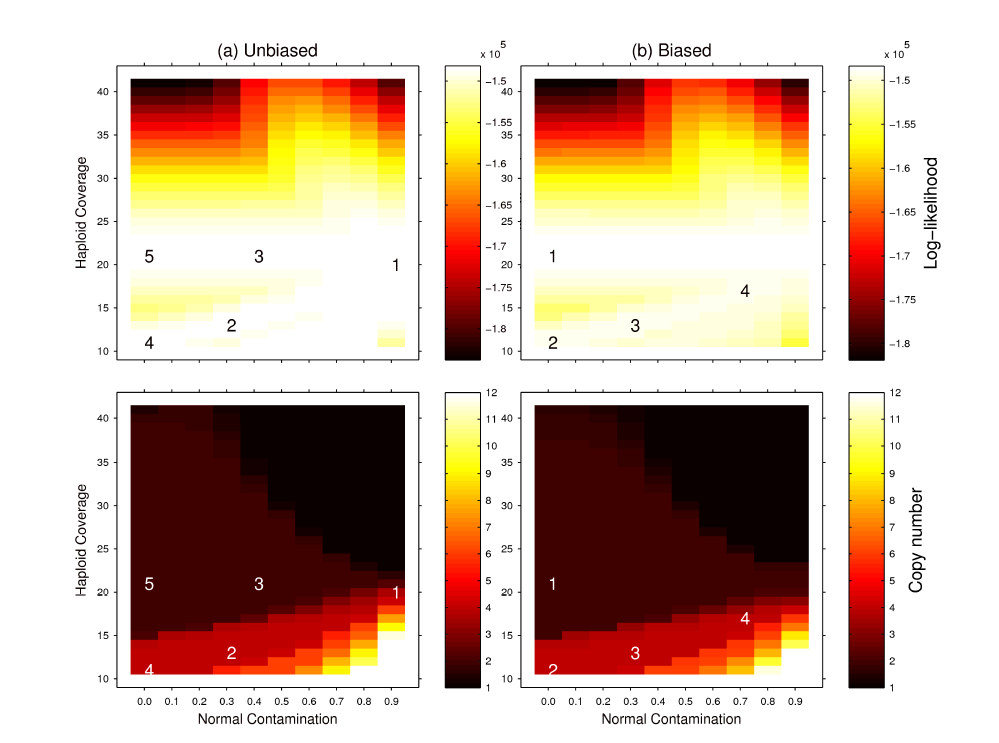
Prior calibration. Prior preferences strongly influence which copy number configuration will be reported to be most probable. When no preference is specified for diploidy (*λ*_3_ = 0) the configuration with highest likelihood has a tetraploid state (left). A suitable calibration for the diploidy parameter can be obtained by selecting a parameter value (*λ*_3_ = 0.05) that makes the diploid state the most probable for tumour samples where the ploidy is known (right).

In OncoSNP-SEQ2, it is possible to calibrate the prior information to *bias* towards certain copy number configurations. For instance, in Figure 8, OncoSNP-SEQ2 identifies five local modes suggesting that there maybe up to five distinct copy number configurations that could plausibly fit the observed data. In an unbiased analysis, where no prior information is supplied, the near-tetraploid copy number configurations are the most probable. However, CLL genomes are generally near-diploid and by adjusting the bias toward diploidy it was possible to force near-diploid solutions to be the most probable. In this case, the diploidy parameter was chosen as the minimal amount necessary to switch the most probable mode from a tetraploid to diploid state and this calibration was used in the subsequent simulations.

## Mathematical Specification

### Prior model

Let 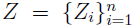 denote the copy number state at *n* SNPs and 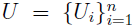 denote the proportion of tumour cells having the corresponding alteration given in *Z* at each SNP. These hidden states are unobserved and form a coupled Markov process such that the transition probabilities have the form:

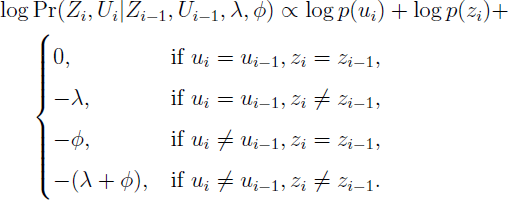

where *p*(*u_i_, z_i_*) denote global priors on the prevalence and copy number states and (*λ, ϕ*) are the transition penalties for changes in *Z* and *U* respectively. Note that in actual implementation we define separate sequences for each chromosome but for brevity of presentation here we assume a single sequence. We use global priors of the form 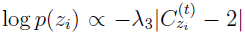 where *C_z_* is the copy number associated with state *z* to bias the solutions towards diploidy when *λ*_3_ *>* 0 and a Beta prior for *p*(*u_i_*).

Furthermore, let 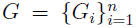 denote the genotypes of the SNP, the state space of *G_i_* depends on *Z_i_*, e.g. if *Z_i_* is a normal copy number two state then *G_i_ ∈ {AA, AB, BB}* or if *Z_i_* is a duplication state then *G_i_ ∈ {AAA, AAB, ABB, BBB}*. Table 2 illustrates genotypes and their respective probabilities for illustrative copy number states.

### Observation model

The observed data is a set 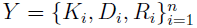 that denotes the alternate allele counts, the total number of reads at each SNP *i* and the average coverage at a set of non-overlapping windows centred on each SNP. For paired normal-tumour data, two observation sequences are observed and unobserved normal and tumour genotypes are coupled in the model. In our simulations, we assume that *D_i_* = *R_i_* and utilise only the total read count overlapping the SNP as a measure of local coverage. However, in general, the full set of sequencing reads can be used.

We model the distribution of the alternate allele count *K_i_* given the total read count *D_i_* is given by a Binomial distribution with density

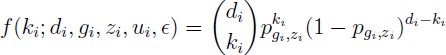

where *p_gi,zi_* = (1 − *ϵ*)*p̂_g_i_,z_i__* + *ϵ*(1 − *p̂_g_i_,z_i__*) is the probability of observing the alternate allele, *ϵ* is the global read error rate and the expected alternate allele probability is given by

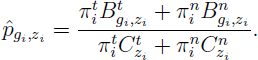

Where 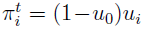 proportion of tumour cells with an alteration, 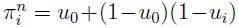 the proportion of cells without the alteration, 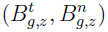 denotes the number of *B* alleles associated with the tumour and normal genotype for the *g*-th genotype state of the *z*-th copy number state and 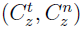 denote the tumour and normal copy numbers.

The distribution of *R_i_* is approximated by a Student-*t* distribution with density

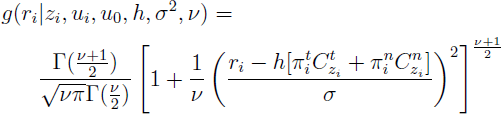

where *h* is the haploid coverage and *σ*^2^ defines the variability of the read coverage. For our simulations we shall assume a sufficiently high sequencing coverage (*>* 60X which is typical in cancers) so that this continuous approximation applies well to discrete data.

### Posterior Inference

We use a grid search method to evaluate identify the most probable joint sequence (*ẑ*, *û*) for different values of baseline coverage *h* and normal cell contamination *u*_0_ using the Viterbi algorithm and evaluate its likelihood *p*(*ẑ*, *û|h, u*_0_, *y, θ*) where *θ* denotes all other model parameters.

From the likelihood grid generated, we identify global maxima and local maxima and report the copy number states and degree of abnormality for each relevant configuration of (*h, u*_0_) and its associated likelihood.

### Multi-scale Copy Number Calling

The *j*-th ranked segmentation is obtained by using the Viterbi algorithm to find the most probable sequence (*z, u*):

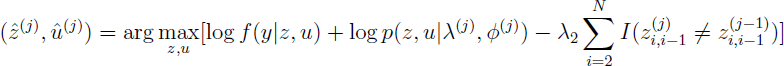

where we use a range of values for *λ* = *ϕ* between 10 and 10^3^ and *λ*_2_ = 1. The final term is a penalty parameter that couples adjacently ranked segmentations to enforce continuity of breakpoints across different segmentations.

### Parameter Settings

By default we set *ν* = 4 for robustness and set 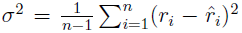 where *ȓ* is a locally smoothed version of *r*. The read error probability is given by *ϵ* = 0.01. These parameters can be learnt from data but results are generally robust with these default parameters. Transition penalties *λ* = 30 and *ϕ* = 30.

### Code availability

The software OncoSNP-SEQ2 is available from: https://sites.google.com/site/oncosnpseq/

## Author's contributions

CY developed methods, analyzed data and wrote the manuscript.

## Acknowledgements

CY acknowledges the support of the UK Medical Research Council through a New Investigator Research Grant (Ref No. MR/L001411/1).

